# Erythroid differentiation dependent interaction of VPS13A with XK at the plasma membrane of K562 cells

**DOI:** 10.1101/2023.08.09.552634

**Authors:** Chase Amos, Peng Xu, Pietro De Camilli

**Affiliations:** Departments of Neuroscience and of Cell Biology, Howard Hughes Medical Institute, Program in Cellular Neuroscience, Neurodegeneration and Repair, Yale University School of Medicine, New Haven, CT 06510; Aligning Science Across Parkinson’s (ASAP) Collaborative Research Network, Chevy Chase, MD 20815

**Keywords:** lipid transfer protein (LTP), endoplasmic reticulum (ER), plasma membrane (PM), membrane, contact

## Abstract

Mutations of the bridge-like lipid transport protein VPS13A and of the lipid scramblase XK result in Chorea Acanthocytosis (ChAc) and McLeod syndrome (MLS) respectively, two similar conditions involving neurodegeneration and deformed erythrocytes (acanthocytes). VPS13A binds XK, suggesting a model in which VPS13A forms a lipid transport bridge between the ER and the plasma membrane (PM) where XK resides. However, studies of VPS13A in HeLa and COS7 cells showed that this protein localizes primarily at contacts of the ER with mitochondria. Overexpression of XK in these cells redistributed VPS13A to the biosynthetic XK pool in the ER but not to PM localized XK. Colocalization of VPS13A with XK at the PM was only observed if overexpressed XK harbored mutations that disengage its VPS13A binding site from an intramolecular interaction. As the acanthocytosis phenotype of ChAc and MLS suggests a role of the two proteins in cells of the erythroid lineage, we explored their localization in K562 cells, which differentiate into erythroblasts upon hemin addition. When tagged VPS13A was overexpressed in hemin treated K562 cells, robust formation of ER-PM contacts positive for VPS13A were observed and their formation was abolished in XK KO cells. ER-PM contacts positive for VPS13A were seldomly observed in undifferentiated K562 cells, in spite of the presence of XK in these cells at concentrations similar to those observed after differentiation. These findings reveal that the interaction of VPS13A with XK at ER-PM contacts requires a permissive state which depends upon cell type and/or functional state of the cell.

## Introduction

Chorea-Acanthocytosis (ChAc) and McLeod syndrome (MLS) are two similar clinical conditions, collectively called neuroacanthocytosis, characterized by degeneration of the brain caudate nucleus, resulting in Huntington’s disease-like neurological manifestations, and presence of abnormal red blood cells (acanthocytes) (Jung *et al*., 2011). ChAc is due to recessive mutations in VPS13A (Rampoldi *et al*., 2001; Ueno *et al*., 2001), an RBG (Repeating Beta-Groove) motif bridge-like lipid transport protein localized at membrane contact sites involving the ER (Kumar *et al*., 2018; Dziurdzik and Conibear, 2021; Leonzino *et al*., 2021; Levine, 2022; Hanna *et al*., 2023). MLS is due to recessive mutations in XK (Ho *et al*., 1994), a lipid scramblase localized in the PM (Adlakha *et al*., 2022; Ryoden *et al*., 2022). Recent studies have shown that the two proteins interact (Urata *et al*., 2019; Park and Neiman, 2020) via the binding of the PH domain of VPS13A to a β-strand hairpin in the 2nd cytosolic loop of XK (Guillen-Samander *et al*., 2022; Park *et al*., 2022) (Fig. 1). Moreover, studies of regulatory T cells have shown that lack of either protein results in a defect of externalization at the PM of PtdSer, a phospholipid normally concentrated in its cytoplasmic leaflet (Ryoden *et al*., 2022).This led to a putative model according to which VPS13A and XK are functional partners: VPS13A would deliver phospholipids including PtdSer from the ER to the PM, while XK would collapse their asymmetric distribution between the two PM leaflets (Adlakha *et al*., 2022; Guillen-Samander *et al*., 2022; Park *et al*., 2022; Ryoden and Nagata, 2022).

**Figure 1.**
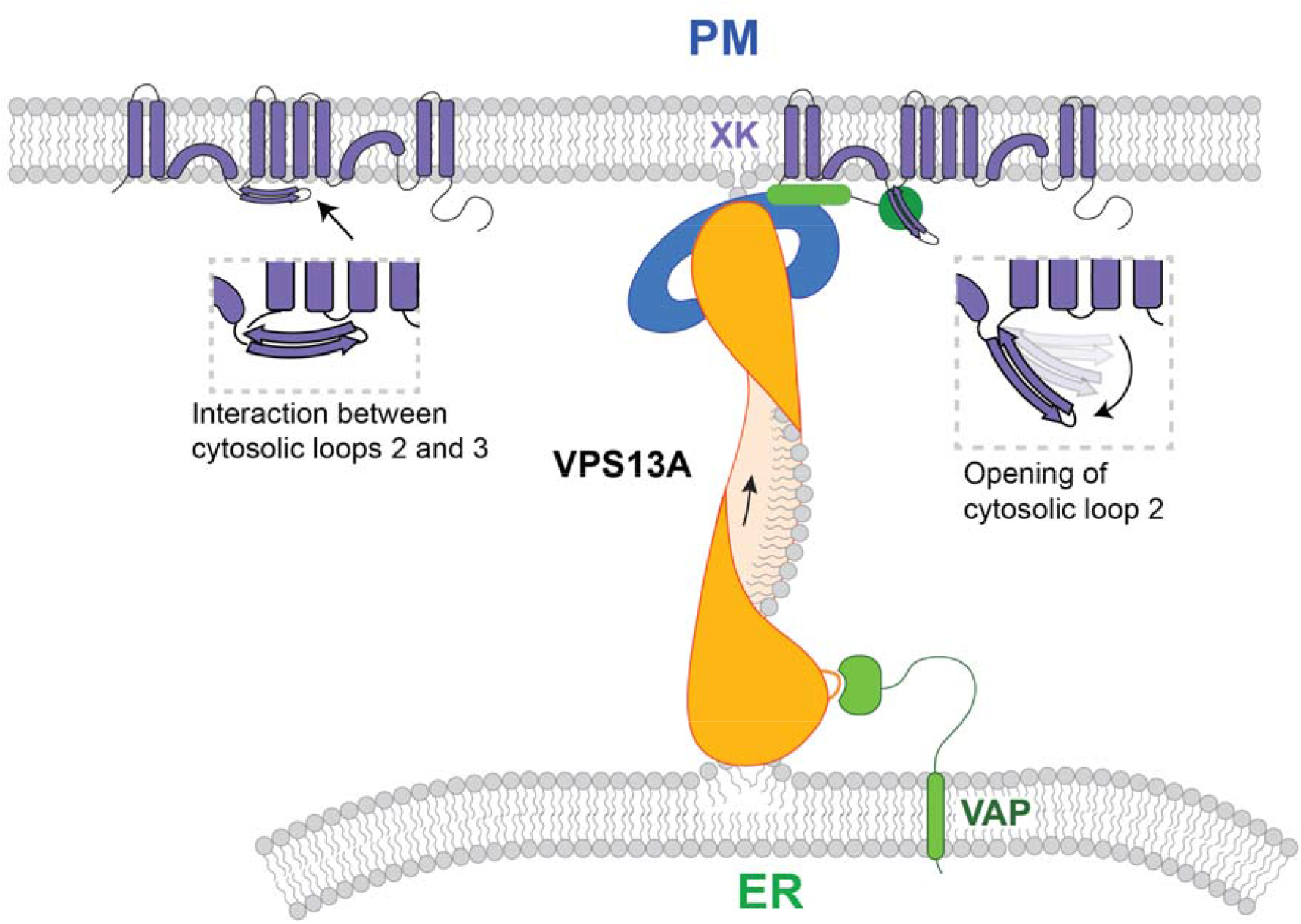
Model of VPS13A interaction with XK at the PM as proposed in Guillen-Samander et al. 2022. VPS13A binds VAP in the ER via an FFAT motif located in its N-terminal region, whereas its C-terminal PH domain binds the second cytosolic loop of XK. When explored in fibroblastic cells by exogenous expression of VPS13A and XK, the interaction of the two proteins at ER-PM contacts occurs only when an intramolecular interaction between loop 2 and loop 3 of XK (magnified in inset) is disrupted by point mutations. (Modified from Guillen-Samander et al., 2022)

However, imaging based studies of tagged VPS13A expressed in commonly used fibroblastic cell lines (COS7 and HeLa cells) had failed to demonstrate an enrichment of VPS13A at ER-PM contact sites, even upon co-expression of WT XK (Guillen-Samander *et al*., 2022). In these cells, VPS13A, when expressed alone, localizes primarily to ER-mitochondria contacts (Kumar *et al*., 2018; Guillen-Samander *et al*., 2022). This interaction occurs via the binding of its FFAT motif containing N-terminal region to the ER protein VAP and of its C-terminal PH domain-containing region to a yet unknown binding site in the outer mitochondrial membrane (Kumar *et al*., 2018; Guillen-Samander *et al*., 2022). Co-expression of WT XK in these cells abolishes the localization of VPS13A to ER-mitochondria contacts, but does not result in its relocation to ER-PM contacts (Park and Neiman, 2020; Guillen-Samander *et al*., 2022). Under these conditions, VPS13A only colocalizes with the biosynthetic pool of XK in the ER, often in focal OSER-like accumulations, while no VPS13A can be found next to the PM pool of XK (Guillen-Samander *et al*., 2022). This relocation is due to binding of the PH domain of VPS13A to ER-localized XK, which outcompetes its binding to mitochondria and likely results in an association of VPS13A with the ER mediated by both its N-terminal and C-terminal regions which bind VAP and XK respectively (see Fig. S3 in Guillen-Samander et al., 2022). While the PH domain of VPS13A can bind XK at the PM, transfected full length VPS13A localizes with XK at the PM, where it also induces a prominent expansion of ER-PM contact sites, only if point mutations are introduced in XK to disrupt a predicted intramolecular interaction between its 2^nd^ and 3^rd^ cytosolic loop (XK^KKR>AAA^ and XK^EYE>AAA^) (Fig. 1) (Guillen-Samander *et al*., 2022). These findings suggested that formation of VPS13A and XK-dependent ER-PM contacts is controlled by regulatory mechanisms that can be bypassed by mutant XK.

We considered the possibility that such mechanisms may be cell type specific. As the acanthocytosis phenotype of ChAc and MLS suggests a special importance of VPS13A and XK in cells of the erythroid lineage, we explored the targeting of VPS13A in a model cell line with properties of such lineage. To this aim we used K562 myelogenous leukemia cells which undergo erythroid differentiation upon treatment with hemin, as shown by elevated hemoglobin content and erythroid lineage-specific transcriptional changes (Baliga *et al*., 1993; Huo *et al*., 2006), including the initial induction of mitophagy as part of organelle clearance (Zhang *et al*., 2009; Fader *et al*., 2016). In these cells we find that, after hemin-induced differentiation, VPS13A is present at ER-PM contacts and that this localization is dependent on endogenous XK, validating the hypothesis that they function together in a disease relevant cell lineage.

## Results

We first investigated the localization of VPS13A in undifferentiated K562 cells. As available antibodies do not detect endogenous VPS13A, we expressed a previously developed plasmid harboring an internal Halo tag, VPS13A^Halo (Kumar *et al*., 2018; Guillen-Samander *et al*., 2022). Similarly to what we had observed in HeLa and COS7 cells, this protein primarily localized at mitochondria, although in a pattern not precisely overlapping with the entire mitochondrial surface, as expected for a selective localization at ER-mitochondria contacts (Fig. 2A). In HeLa and COS7 cells, VPS13A^Halo also localizes at ER-lipid droplet contacts, but no lipid droplets were present in our K562 cells. Next, we investigated the localization of VPS13A after hemin-induced differentiation. Consistent with previous reports (Baliga *et al*., 1993), incubation of cells in the presence of hemin for 2 days induced expression of hemoglobin with variable efficiency, as detected by a diaminobenzidine-based cytochemical reaction, confirming differentiation toward the erythroid lineage (Fig. 3). In striking contrast to what we had observed in undifferentiated cells, expression of VPS13A^Halo in hemin-treated cells resulted in its localization not only at mitochondria but also at the PM in a large fraction of the cells (Fig. 2A-D). This localization had the expected appearance of ER-PM contacts, as seen both in equatorial (positive segments along the PM) (Fig. 2B) and basal planes (patches parallel to the substrate) (Fig. 2C). Moreover, expression of VPS13A^Halo and of the ER marker GFP-Sec61β showed that cortical patches of VPS13A were connected to ER tubules (Fig. 2E). The abundance and extent of these contacts was somewhat variable, most likely reflecting the variable efficiency of differentiation.

**Figure 2.**
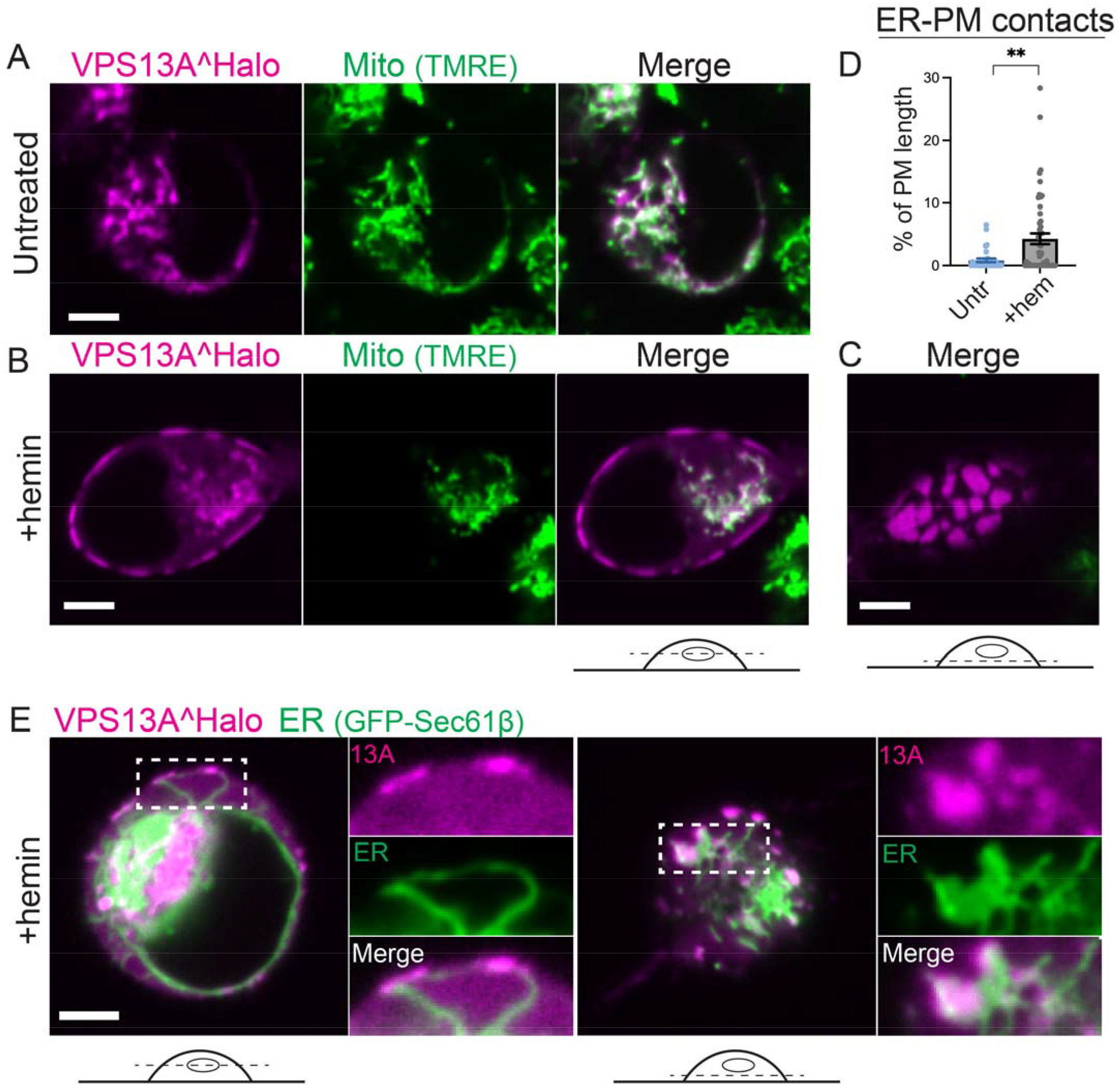
Overexpressed VPS13A is recruited to ER-PM contacts of K562 cells following their differentiation to the erythroid lineage by hemin. (A-B) Equatorial views of live K562 cells expressing VPS13A^Halo and untreated (A) or treated (B) with 30 μM hemin for 2 days and imaged after a brief incubation with TMRE, a live mitochondrial marker. Panel C shows the basal plane of the cell shown in the left panels revealing “en face” views of large ER-PM contacts. (D) Quantification of the fraction of PM profiles with VPS13A enrichment (D) (n = 31 untreated cells, 52 hemin-treated cells). (E) Equatorial (left) and basal (right) views of hemin treated K562 cells co-expressing VPS13A^Halo and GFP-Sec61β. White rectangles outline areas shown at high magnification at the right of the main fields.** indicates p-value<0.01. Scalebars indicate 5 μm.

**Figure 3.**
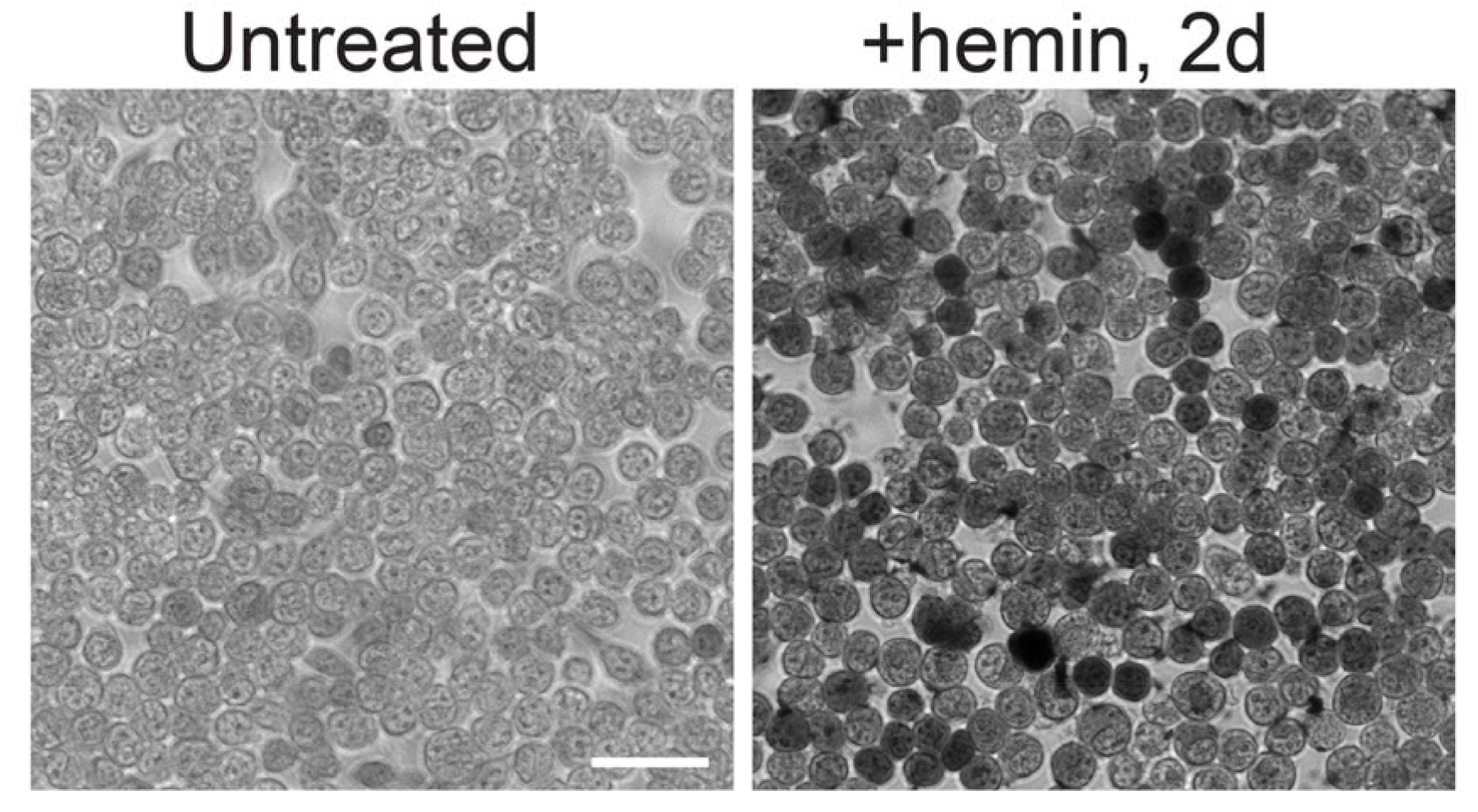
Differentiation of K562 cells to the erythroid lineage upon addition of hemin for 2 days. Brightfield image of cells untreated (left) or treated (right) with 30 μM hemin for 2 days and processed through a diaminobenzidine/H_2_O_2_ based cytochemical reaction to reveal presence of hemoglobin (dark gray). Scalebar indicates 50 μm.

We next examined the impact of expressing exogenous XK on these localizations. As previously observed in COS7 cells (Guillen-Samander *et al*., 2022), co-expression of GFP-XK with VPS13A^Halo in non-differentiated K562 cells abolished the localization of VPS13A^Halo at mitochondria and resulted in its diffuse distribution throughout the cell, most likely reflecting colocalization with a pool of XK in the ER (Fig. 4A). While a pool of GFP-XK^WT^ was observed at the PM in these cells, VPS13A^Halo did not colocalize with this pool. In contrast, in hemin-treated cells expressing GFP-XK^WT^, VPS13A was concentrated at PM patches (Fig. 4A, C). Moreover, and in agreement with what we had observed in COS7 cells, co-expression of VPS13A^Halo with a GFP-XK construct harboring mutations in its 2^nd^ cytosolic loop (K105A/K106A/R107A, referred to as XK^KKR>AAA^) which disrupts its interaction with the 3^rd^ loop (Guillen-Samander *et al*., 2022), accumulated at ER-PM contacts regardless of differentiation status (Fig. 4B-C).

**Figure 4.**
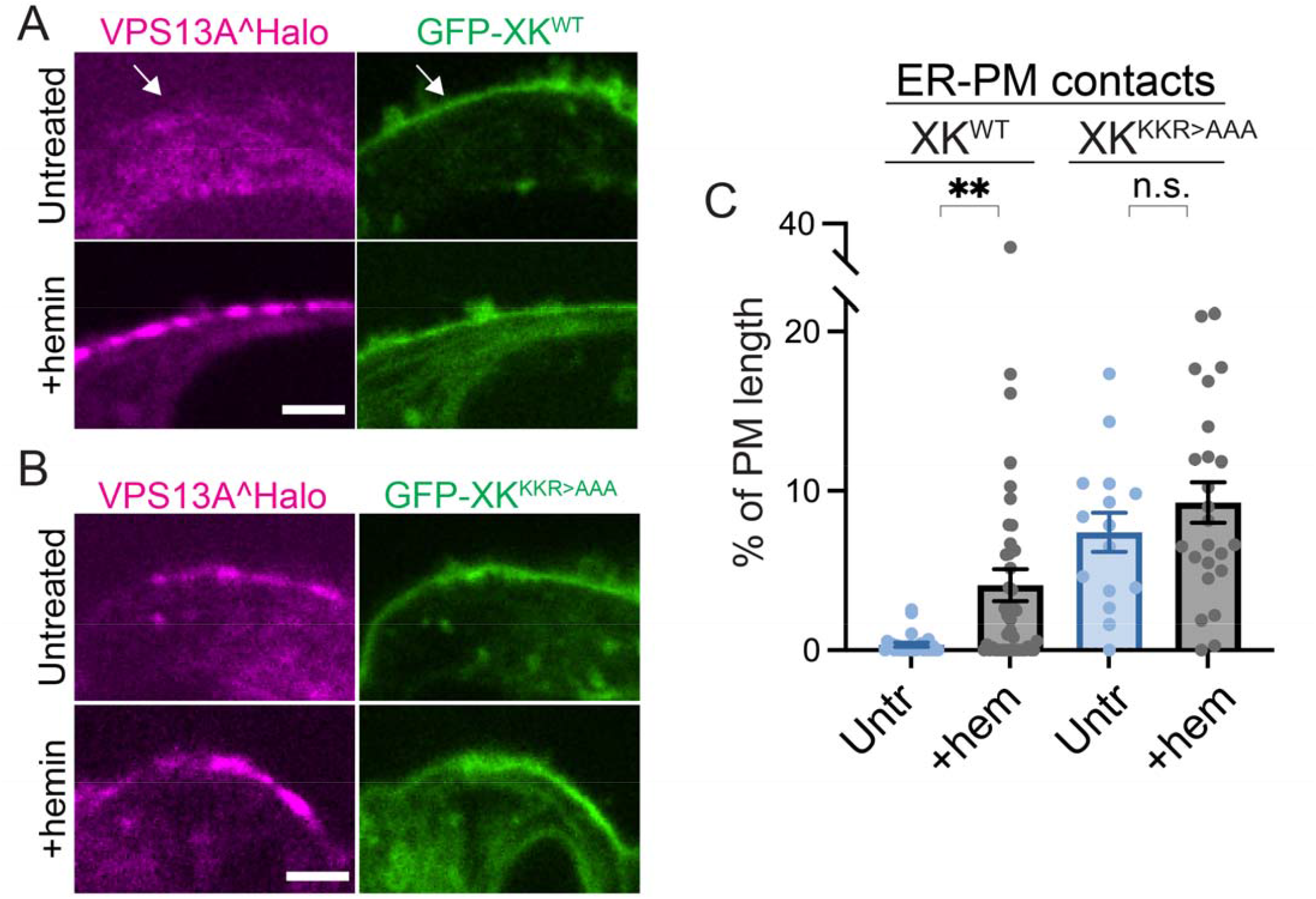
Expression of exogenous XK^KKR>AAA^, but not of exogenous XK^WT^, recruits VPS13A to ER-PM contacts also in undifferentiated K562 cells. (A-B) Co-expression of GFP-XK^WT^ (A) or GFP-XK^KKR>AAA^ (B) and VPS13A^Halo in untreated or hemin-treated cells, with a magnified view of the PM shown. (C) Quantification of fraction of PM with VPS13A (n=26 untreated XK^WT^ cells, 46 hemin-treated XK^WT^ cells, 15 untreated XK^KKR>AAA^ cells, 24 hemin-treated XK^KKR>AAA^ cells). ** indicates p-value<0.01, n.s. p-value>0.05. Scalebars indicate 5 μm.

The most plausible interpretation of these findings is that the recruitment of VPS13A to the PM in differentiated cells is due to hemin-dependent expression of endogenous XK. To address this possibility, we analyzed expression of XK in K562 cells and we generated K562 cells lacking XK by CRISPR/Cas9 gene editing. Western blotting analysis of lysates surprisingly revealed that a band with the expected mobility of XK (Ryoden *et al*., 2022), and absent in XK KO K562 cells, was already enriched in non-differentiated cells (Fig. 5A, lane 2). Total XK did not increase, and in fact showed a mild, non-significant decrease after hemin (Fig. 5A-B), potentially due to early stages of organelle clearance during differentiation towards erythrocytes. However, the recruitment of VPS13A^Halo to the PM after hemin was abolished in KO cells, while the localization of VPS13A at mitochondria persisted, confirming the role of XK in its PM recruitment (Fig. 5C-E). Note in Fig. 5A the low expression of XK in COS7 cells relative to K562 cells, demonstrating cell type differences in the expression levels of XK. These findings demonstrate that presence of endogenous XK is not sufficient to mediate VPS13A targeting to the PM, pointing to the occurrence of regulatory mechanisms that correlate with erythroid differentiation.

**Figure 5.**
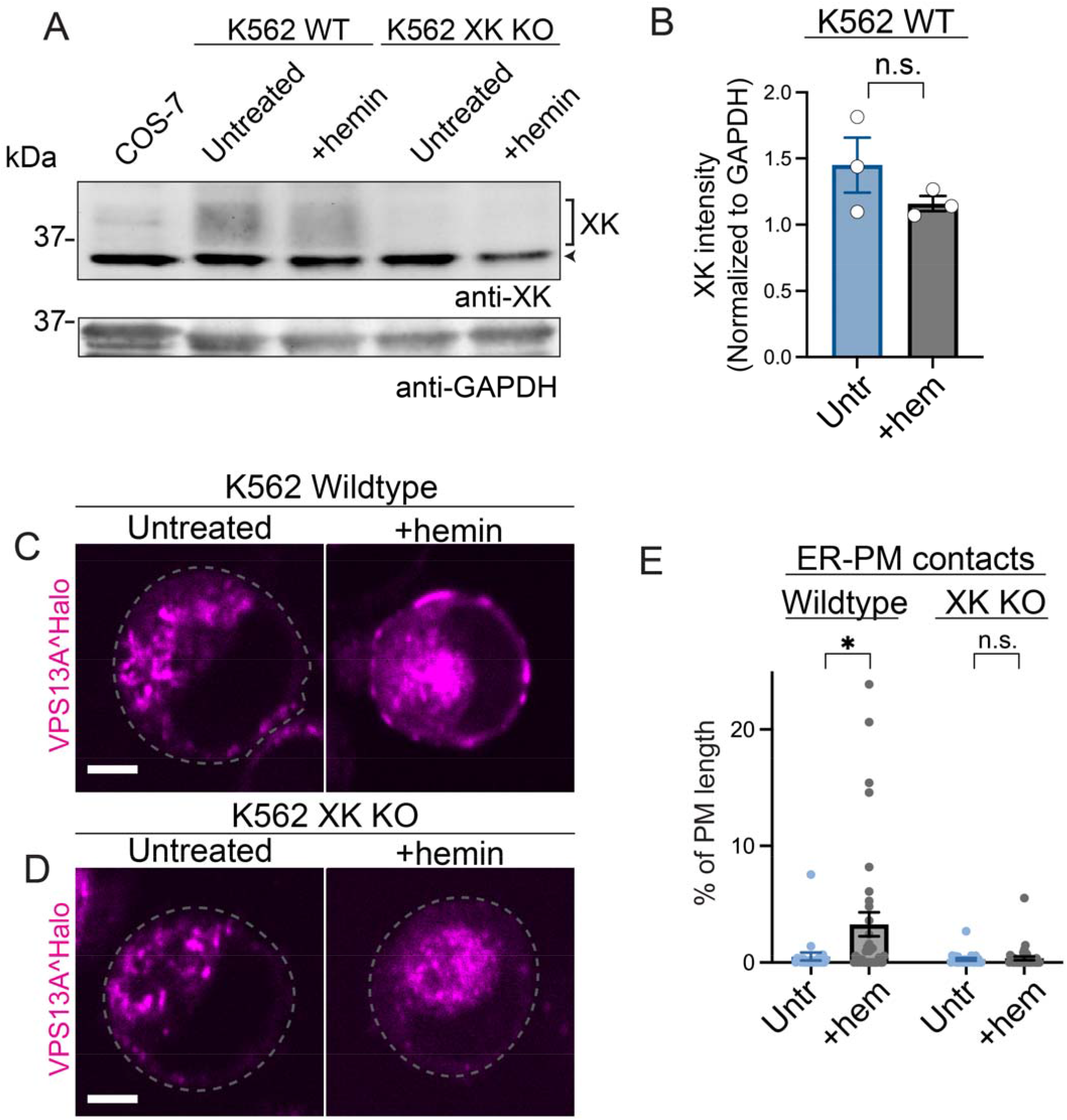
Recruitment of overexpressed VPS13A to the PM of erythroid cells depends on XK. (A-B) Western blot of lysates from COS-7 cells and wildtype or XK KO K562 cells untreated or treated with hemin for 2 days (A), with quantification of XK intensities normalized to GAPDH input from western blots in (B) (n=3 untreated lysates, 3 hemin-treated lysates). An arrowhead indicates a non-specific band. (C-E) Wildtype (C) and XK KO (D) cells untreated or treated with hemin were transfected with VPS13A^Halo. Stippled lines indicate the cell profile in images where such profiled is not visible. The fraction of the PM profiles occupied by VPS13A is shown in (E) (n=23 wildtype untreated cells, 35 wildtype hemin-treated cells, 24 XK KO untreated cells, 36 XK KO hemin-treated cells). * indicates p-value < 0.05, n.s. p-value>0.05 by paired (B) or unpaired (E) Student’s t-test. Scalebars indicate 5 μm.

## Discussion

Our findings prove that in erythroblast model cells, i.e. cells of a lineage that undergoes dysfunction upon loss of either VPS13A or XK, overexpressed VPS13A localizes at ER-PM contact sites dependently on XK. We also find that presence of endogenous XK is not sufficient to result in their colocalization at ER-PM contacts. While level of endogenous XK in K562 cells are similar before and after hemin-induced differentiation, such localization only largely occurs in hemin differentiated cells. Thus, differentiation of K562 toward the erythrocyte lineage correlates with a cellular status that is permissive for the binding of full length VPS13A to WT XK at the PM.

These mechanisms likely impact the intramolecular interaction between the 2^nd^ and 3^rd^ intracellular cytosolic loop of XK, as both in COS7 cells, where XK expression is comparatively low, and in undifferentiated K562 cells, which express robust levels of endogenous XK, this intramolecular interaction must be disrupted to allow full length VPS13A binding at the PM. Based on studies of the XK paralogue XKR8, the 3^rd^ intracellular loop of XK is likely to play a regulatory role on the scrambling activity of XK (Sakuragi *et al*., 2021). Thus, the involvement of this loop of XK in the regulation of VPS13A binding may reflect an interplay between VPS13A binding and the regulation of lipid scrambling. Why the binding of full length VPS13A to endogenous XK at the PM, but not to XK in the ER, is affected by this erythroid differentiation-dependent switch in K562 cells, remains an open question.

In regulatory T cells, the partnership of VPS13A with XK is thought to be important for the externalization of PtdSer. Interestingly, *Drosophila* mutants lacking the VPS13A/VPS13C orthologue show a defect in the removal of cell corpses (Faber *et al*., 2020), a process that typically requires PtdSer exposure. Moreover, in erythroid differentiation, PtdSer externalization plays a role during enucleation of erythroblasts when macrophages surround and then engulf the portion of the cell containing the nucleus (Moras *et al*., 2017). Thus, one could expect a defect in this process in cells lacking VPS13A or XK. However, so far, the main blood cell defect observed in ChAc and MLS is the presence of acanthocytes, i.e. a defect of mature red cell morphology, rather than of red cell enucleation. In view of the scrambling activity of XK, it is of interest that acanthocytes can be the result of abnormal bilayer asymmetry leading to buckling of the membrane (Sheetz and Singer, 1974; Redman *et al*., 1989).

VPS13A is still present in the PMs of mature red cells, where its levels are drastically reduced in the absence of XK, suggesting that its interaction with XK stabilizes it (Urata *et al*., 2019). In fact, lack of VPS13A in red cell membranes has been used to diagnose ChAc (Peikert *et al*., 2023). Thus, as erythroblasts undergo full maturation to erythrocytes, VPS13A remains associated with XK in spite of the disappearance of the ER. A major outstanding open question is what may be the function of this protein which is thought to transport lipids form the ER to the PM, once the ER is no longer present. One possibility is that a regulatory function of VPS13A on the lipid scrambling activity of XK, irrespective of the lipid transport properties of VPS13A, may be preserved in mature red cells.

## Methods

### Plasmids

The plasmid encoding VPS13A^Halo (Kumar *et al*., 2018) (RRID:Addgene_118759) and GFP-XK(KKR>AAA mutant) (Guillen-Samander *et al*., 2022) were generated in our lab. GFP-XK was obtained from Genscript. GFP-Sec61β was a gift from T. Rapoport (Harvard University, Cambridge, MA) and mito-BFP from G. Voeltz (University of Colorado Boulder, Boulder) (RRID:Addgene_49151).

### Cell culture and imaging

K562 cells (gift of Patrick Gallagher, Yale University) were cultured at 37°C in 5% CO2, in RPMI 1640 supplemented with FBS, GlutaMAX, non-essential amino acids, and sodium pyruvate (Gibco). COS-7 cells (ATCC Cat# CRL-1651, RRID:CVCL_0224) used for lysates were cultured in DMEM medium (Gibco) with the same supplements. Prior to transfections, cells were maintained with penicillin/streptomycin (Gibco) and plasmocin (InvivoGen). For transections, cells were plated on fibronectin (Sigma Aldrich) coated 35 mm imaging dishes (Mattek) in the same medium but without antibiotics and incubated overnight. Cells were then rinsed with medium to remove non adhering cells and cell debris, and new medium was added containing the plasmids, FuGene 4K Transfection Reagent (Promega) as well as hemin in a subset of dishes. Hemin (Sigma Aldrich) dissolved in DMSO was added at a final concentration of 30 μM. After overnight incubation with hemin and transfection reagents, the medium was replaced with new medium containing the same factors for an additional overnight incubation. After 2 days following transfection and hemin treatment, cells were incubated with Halo ligands (gift of L. Lavis, Janelia Research Campus) for 90min and rinsed with RPMI media. Immediately prior to imaging, live cells were stained with TMRE (Cayman Chemical, TMRE Mitochondrial Membrane Potential Assay Kit). Imaging was performed at 37°C, 5% CO2 on an Andor Dragonfly (PlanApo 63x objective, 1.4 numerical aperture) microscope equipped with a Zyla cMOS camera. A gaussian blur is applied to some of the images presented.

For diaminobenzidine staining, a final concentration of 0.2% 3,3’-diaminobenzidine tetrahydrochloride hydrate and 0.03% hydrogen peroxide (Sigma Aldrich) was combined in PBS immediately prior to staining. K562 cells on Mattek dishes were incubated with the diaminobenzidine/hydrogen peroxide mixture for 10 min at 37°C, 5% CO2, washed with PBS, and the brightfield view was imaged with a ZOE Fluorescence Cell Imager (Bio-Rad).

Detailed protocol for cell culture, transfection and imaging: https://dx.doi.org/10.17504/protocols.io.e6nvwdk4dlmk/v1 (Private link for reviewers: https://www.protocols.io/private/3DB6421E2FD011EE96830A58A9FEAC02 to be removed before publication.)

### Image analysis

The fraction of PM occupied by VPS13A^Halo was quantified using FIJI (RRID:SCR_002285). Due to the close proximity of mitochondria to the cell edge, a negative mask generated from TMRE staining or mito-BFP overexpression was used to subtract VPS13A^Halo signal originating from the mitochondria. The mitochondria-subtracted VPS13A channel was then thresholded to generate a binary mask of signal at the PM. After tracing the cell edge, the percentage of the PM length occupied by the binary VPS13A mask was measured.

Detailed protocol for image analysis of PM contacts: https://dx.doi.org/10.17504/protocols.io.n2bvj364plk5/v1 (Private link for reviewers: https://www.protocols.io/private/2752B9812FD611EE9BA00A58A9FEAC02 to be removed before publication.)

### Gene editing

The knockout of XK was performed using a previously published CRISPR/Cas9 plasmid with a gRNA targeting exon 1 of XK (Guillen-Samander *et al*., 2022). The backbone was plasmid px459 (Addgene #62988) with the following XK gRNA sequence CCGTTGTCTCGGCCACGAACAGG (Guillen-Samander *et al*., 2022). K562 cells were transfected with this plasmid using the SF Cell Line 4D-Nucleofector kit and subsequently selected with 1 μg/mL puromycin prior to single cell cloning. Sequencing of the gRNA-targeted exon 1 was performed by PCR amplification (Promega GoTaq polymerase) with 5’-GGTTTGGGGCTGGGCAT-3’ and 5’-AGGTGCAGCAGCAGTACG-3’ primers (Guillen-Samander *et al*., 2022) and cloning with the TOPO TA Cloning Kit (Invitrogen).

Previously published detailed protocol for CRISPR/Cas9 editing of mammalian cells: https://dx.doi.org/10.17504/protocols.io.5jyl85x89l2w/v1.

### Western blotting

K562 and COS-7 cells were lysed in 2% SDS and cleared by centrifugation at 13,300 RPM for 10 min after sonication. Lysates were mixed with a concentrated solution of SDS loading buffer to reach a final concentration of 50 mM Tris pH 6.8, 2% SDS, 0.1% Bromophenol blue, 10% glycerol, 1% beta-mercaptoethanol, heated for 10 min at 95°C, and loaded on 4-12% Tris Glycine gels (Invitrogen). Gels were transferred to nitrocellulose membranes (Biorad) overnight at 30 V, 4°C. After blocking with 5% BSA (Sigma Aldrich) in 1x TBS-T (Santa Cruz, Tris Buffered Saline with Tween) for 1 hour at room temperature, membranes were incubated overnight at 4°C with anti-XK (Sigma-Aldrich Cat# HPA019036, RRID:AB_1846286) and anti-GAPDH (EnCor Biotechnology Cat# MCA-1D4, RRID:AB_2107599 in Fig. 5A; Thermo Fisher Scientific Cat# MA5-15738, RRID:AB_10977387 in Fig. 5B) in 5% BSA in TBS-T. After three washes with TBS-T, membranes were incubated for 1 hour at room temperature with LI-COR IRDye 680CW or 800CW secondary antibodies in 5% BSA in TBS-T, washed three times with TBS-T, and visualized with a Licor Odyssey Infrared Imager.

Detailed protocol for western blotting: https://dx.doi.org/10.17504/protocols.io.3byl4qb6zvo5/v1

(Private link for reviewers: https://www.protocols.io/private/2041DA122FD111EE96830A58A9FEAC02 to be removed before publication.)

## Statistical analysis

Quantifications were analyzed using GraphPad Prism (RRID:SCR_002798). Unless otherwise indicated, all statistical analyses indicated are from an unpaired Student’s t-test and graphs indicate mean ± S.E.M.

## Data availability

All primary data associated with each figure has been deposited in the Zenodo repository and will be available upon publication using the following link: https://doi.org/10.5281/zenodo.8200336/.

## Acknowledgments

We thank Andres Guillén-Samander, Hanieh Falahati, Ben Johnson, and Sydney Cason for discussion. This work was supported in part by NIH grants NS36251 and DA018343 and by the Parkinson Foundation (PF-RCE-1946). The study is funded by the joint efforts of The Michael J. Fox Foundation for Parkinson’s Research (MJFF) and the Aligning Science Across Parkinson’s (ASAP) initiative. MJFF administers the grant ASAP-000580 on behalf of ASAP and itself. For the purpose of open access, the author has applied a CC-BY public copyright license to the Author Accepted Manuscript (AAM) version arising from this submission.

## Notes

### Competing Interest Statement

The authors have declared no competing interest.

